# A single main-chain hydrogen bond required to keep GABA_A_ receptors closed

**DOI:** 10.1101/2024.12.22.629966

**Authors:** Cecilia M. Borghese, Jason D. Galpin, Samuel Eriksson Lidbrink, Yuxuan Zhuang, Netrang G. Desai, Rebecca J. Howard, Erik Lindahl, Christopher A. Ahern, Marcel P. Goldschen-Ohm

## Abstract

GABA_A_ receptors are the primary inhibitory neurotransmitter receptors throughout the central nervous system. Despite significant progress understanding their three-dimensional structure, a critical gap remains in determining the molecular basis for channel gating. We recently identified M2-M3 linker mutations that suggest linker flexibility has asymmetric subunit-specific correlations with channel opening. Here we use non-canonical amino acids (ncAAs) to investigate the role of main-chain H-hydrogen bonds (H-bonds) that may stabilize the M2-M3 linkers. We show that a single main-chain H-bond within the β2 subunit M2-M3 linker inhibits pore opening and is required to keep the unliganded channel closed. Furthermore, breaking this H-bond during channel opening accounts for approximately one third of the activation energy derived from GABA binding. In contrast, the analogous H-bond in the α1 subunit has no effect on gating. Our observations suggest that channel opening involves state-dependent breakage/disruption of a specific main-chain H-bond within the β2 subunit M2-M3 linker.

## Introduction

Main-chain H-bonds are essential for proteins to fold into helix and sheet secondary structures. Despite their obvious role in structure, their mechanistic contribution to the function of mature proteins is typically unknown, especially in less ordered loops where their importance is often not structurally obvious. The relative lack of functional measures that probe main-chain H-bonds is largely due to the fact that conventional site-directed mutagenesis only allows swapping side-chains but does not alter the composition of the main-chain. Nonetheless, there exist well-established methods for introducing ncAAs with altered main-chain chemistry ^1,2^. In particular, α-hydroxy acids enable amide-to-ester substitutions in the main-chain that ablate H-bonds with the amide nitrogen while leaving the side-chain unchanged ^1,3^. Effectively, this enables targeted elimination of specific main-chain H-bonds to test their contribution to protein function.

Although largely under-utilized for ion channels, incorporation of α-hydroxy acids has revealed important roles for main-chain H-bonds in ion selectivity of acid-sensing (ASIC) and chloride (CLC) channels ^4,5^, voltage-sensitivity of Shaker potassium channels ^6^, gating dynamics for inwardly rectifying potassium (Kir) channels ^7,8^, both ligand binding and channel gating in nACh receptors ^9–15^, and stability of the open state in the prokaryotic homolog GLIC ^16^. Here, we determine the mechanistic contribution of a main-chain H-bond within an important gating loop of a typical synaptic γ-aminobutyric acid type A receptor (GABA_A_R).

GABA_A_Rs are members of the cys-loop superfamily of pentameric ligand-gated ion channels (pLGICs) including glycine (Gly), nicotinic acetylcholine (nACh), and serotonin (5-HT3) receptors ^17^. As the primary mediators of inhibitory synaptic signaling throughout the central nervous system, it is perhaps not surprising that genetic mutations causing GABA_A_R dysfunction are related to a broad spectrum of human disorders such as epilepsy, neurodevelopment and intellectual disability, autism spectrum disorder, schizophrenia, and depression. They are also important drug targets for anxiolytics, anticonvulsants, antidepressants, and anesthetics which modulate channel activity, and thereby inhibitory tone in neural circuits.

GABA_A_Rs are comprised of subtype-specific combinations of homologous but nonidentical subunits (α1-6, β1-3, γ1-3, δ, ε, ρε, 8, π1-3) that together form a central chloride-conducting pore through the plasma membrane ^18–20^. Cryo-EM structures of common synaptic subtypes α1β2/3γ2 GABA_A_Rs have been enormously useful for mechanistic inference and prediction (**Figure 1A**) ^19–29^. Agonists (e.g., the neurotransmitter GABA) bind to two sites in the extracellular domain (ECD) at β/α inter-subunit interfaces, where they promote opening of the ion pore in the transmembrane domain (TMD). Other distinct sites mediate allosteric modulation by anxiolytics (e.g., diazepam), anticonvulsants, antidepressants, and anesthetics. Loops at the ECD-TMD coupling interface including the M2-M3 linkers are crucial for transducing the chemical energy from agonist binding to pore gating (**Figure 1B**) ^30–33^. However, the detailed interactions mediating this coupling remain only poorly understood. Comparison of structural snapshots in inactive (e.g., antagonist-bound) and activated/desensitized (e.g., GABA-bound) conformations suggest a prominent motion during agonist-activation is a radial expansion of the M2-M3 linkers which sequentially follow the pore-lining M2 helices ^23,34,35^.

**Figure 1.**
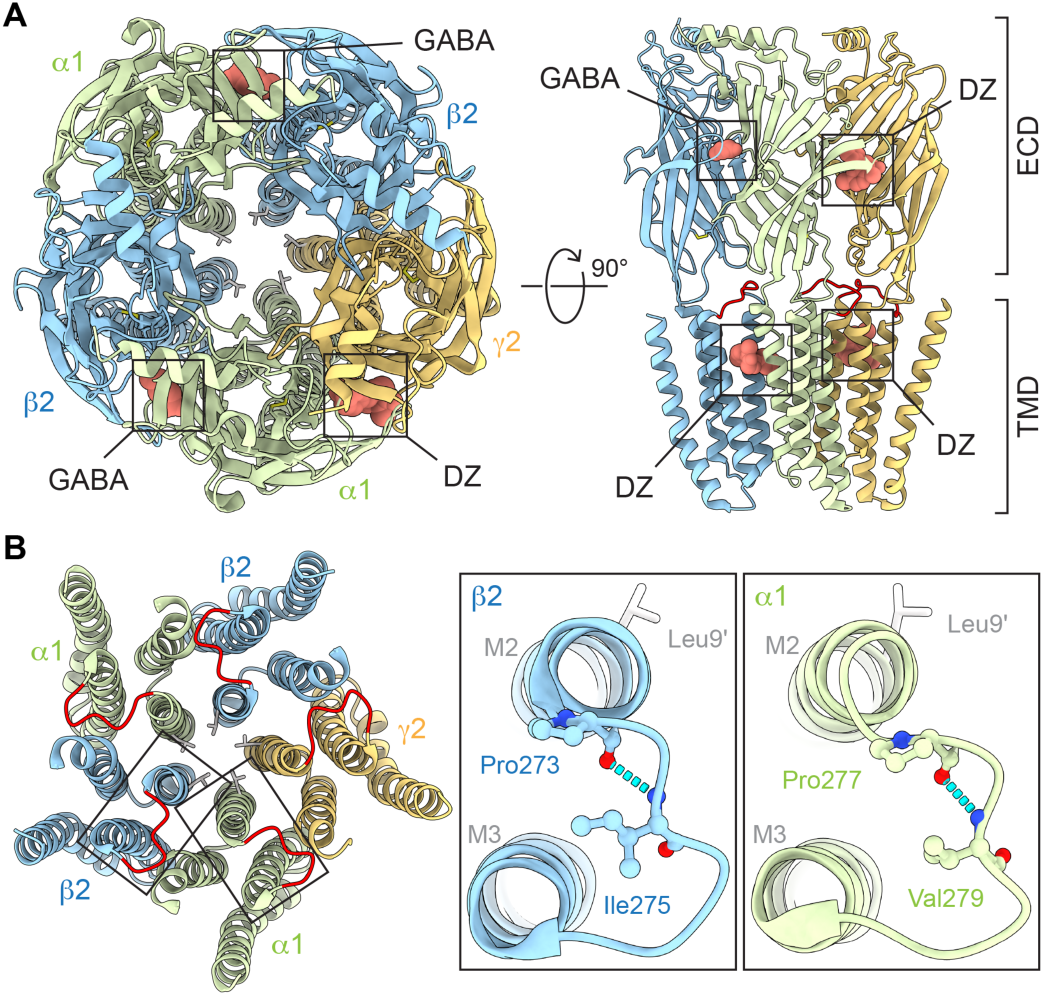
Predicted main-chain H-bonds within M2-M3 linkers. **(A)** Cryo-EM map (PDB 6X3X) of human α1β2γ2 GABA_A_R with bound GABA and the positive modulator diazepam (DZ) viewed from the top (left) and side (right). M2-M3 linkers colored red on right. **(B)** Left: Top view of TMD only for antagonist-bound structure (PDB 6X3S). M2-M3 linkers colored red and 9′ pore gate leucines shown as sticks. Right: M2-M3 linkers for boxed regions on left. Predicted main-chain H-bonds α1(Val279:N-Pro277:O) and β2(Ile275:N-Pro273:O) indicated with cyan dashed segments and involved residues shown as ball and stick. Rat residue numbering shown, which is offset by one from human numbering in α1 (Figure S1). A similar H-bond is predicted in the γ2 subunit (not shown). Structure visualizations with ChimeraX ^43^.

Here, we follow up on our previous observations that alanine substitutions predicted to increase M2-M3 linker flexibility have highly asymmetric subunit-specific effects on gating and diazepam modulation ^36,37^. Structures indicate a potential main-chain H-bond within the M2-M3 linker that could stabilize a crimp or shallow hairpin in the linker backbone predicted to limit linker flexibility (**Figure 1B**). We show that this H-bond in the β2 subunit is a crucial component of the channel gating mechanism, whereas it has no functional role in the α1 subunit despite high structural homology.

## Results

### Main-chain H-bond elimination with ncAAs

To eliminate the putative main-chain H-bonds α1(Val279:N-Pro277:O) or β2(Ile275:N-Pro273:O) within the M2-M3 linkers (**Figure 1B**), either α1(Val279) or β2(Ile275) was substituted with the amber stop codon (TAG), denoted α1(Val279*) or β2(Ile275*). We then used *in vivo* nonsense suppression to introduce the cognate α-hydroxy acid (Vah or Iah) at the cRNA UAG site as previously described ^6^. The amide-to-ester substitutions α1(Val279Vah) or β2(Ile275Iah) effectively eliminate any main-chain H-bonding between the amide nitrogen of α1(Val279) or β2(Ile275) and the carboxylate oxygen of α1(Pro277) or β2(Pro273), respectively, without altering the side-chains of these residues (**Figures 1B**, **2A**). Briefly, Xenopus laevis oocytes were coinjected with cRNA for α1, β2, and γ2 subunits (or cRNA for specific UAG mutants) and an orthogonal pyrrolysine tRNA (a natural UAG suppressing tRNA) that has been chemoenzymatically ligated with either *i*) the cognate α-hydroxy acid (the test case), *ii*) the wild-type (WT) amino acid (should recapitulate WT behavior), or *iii*) nothing (blank full-length tRNA; should not express GABA_A_R due to early termination at the introduced stop codon) (**Figure 2B**). For each batch of oocytes, current responses to pulses of the pore blocker picrotoxin (PTX) and a series of GABA concentrations were recorded from oocytes in all three conditions (**Figure 2C**). Reliable nonsense suppression incorporation of residues with little to no read-through is evident from the lack of GABA_A_R currents from oocytes in condition *iii* as compared to conditions *i* and *ii* (**Figure 2D**) and the recapitulation of WT behavior in condition *ii* (**Figure 3**).

**Figure 2.**
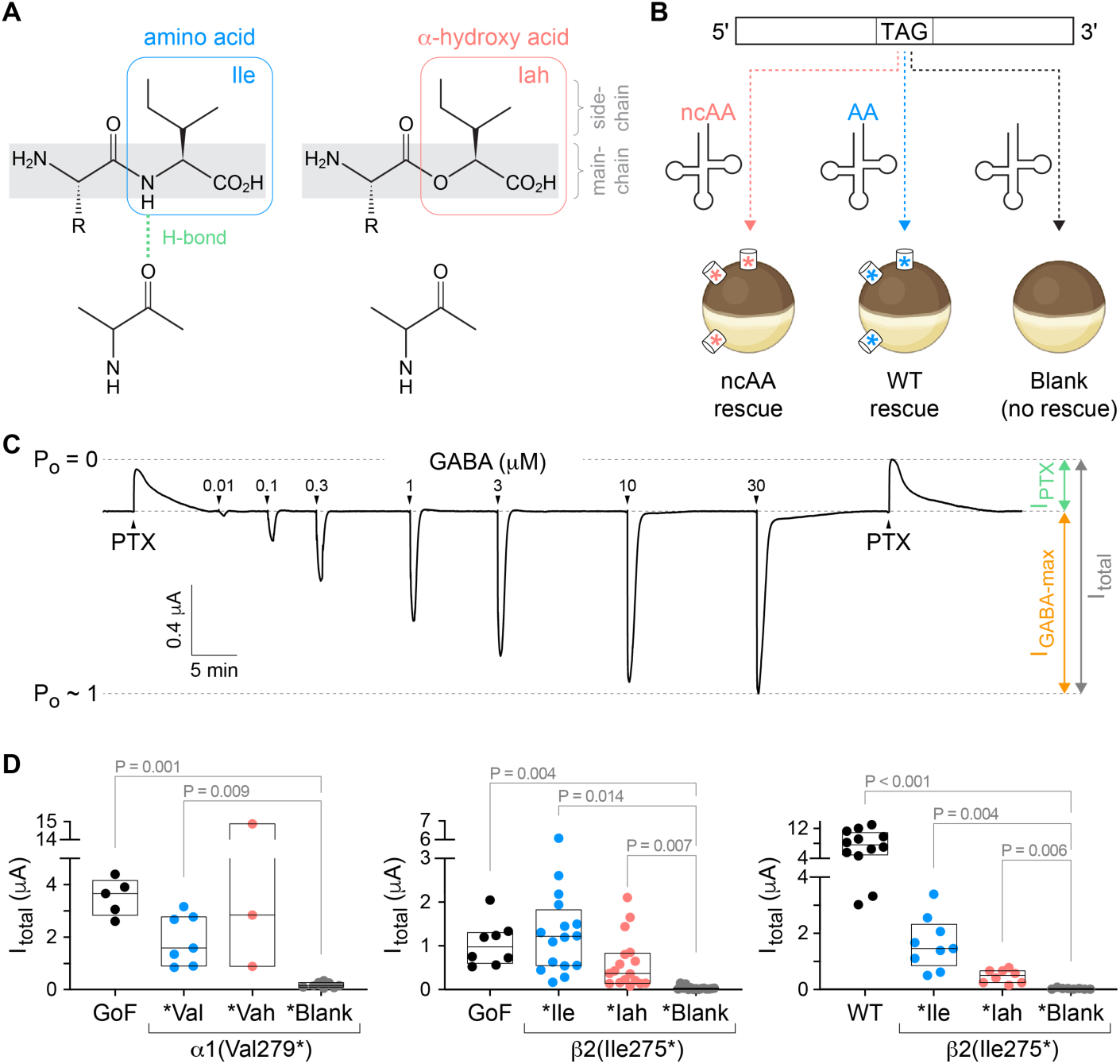
Nonsense suppression incorporation of α-hydroxy acids to eliminate specific main-chain H-bonds. **(A)** Depiction of a main-chain H-bond with the amide NH of an isoleucine (Ile) amino acid residue and its ablation upon substitution with its cognate α-hydroxy acid (Iah). **(B)** Oocytes are coinjected with cRNA for GABA_A_R subunits with a TAG stop codon at the site of interest along with tRNA ligated to either the wild-type amino acid (AA), its cognate α-hydroxy acid (ncAA), or nothing (blank). The result is expression of wild-type channels, channels incorporating the α-hydroxy acid at the TAG site, or no channels due to truncation at the TAG site, respectively. **(C)** Current trace for α1(Leu9′Thr)β2ψ2 gain-of-function (GoF) receptors. Arrows indicate approximate onset of pulses of either 1 mM PTX or increasing concentrations of GABA. Open probability (P_o_) estimated by normalizing from the zero-current baseline in PTX to the maximal current elicited with GABA. **(D)** Total current per oocyte as illustrated in panel c suggests reliable nonsense suppression incorporation of AA and ncAA with little to no read-through (i.e., relative lack of current for blank) in both GoF and WT backgrounds. Box plots show median and interquartile intervals. P-values < 0.05 for Brown-Forsythe ANOVA with posthoc Dunnett’s T3 test show. See Table S1 for summary statistics and number of oocytes.

**Figure 3.**
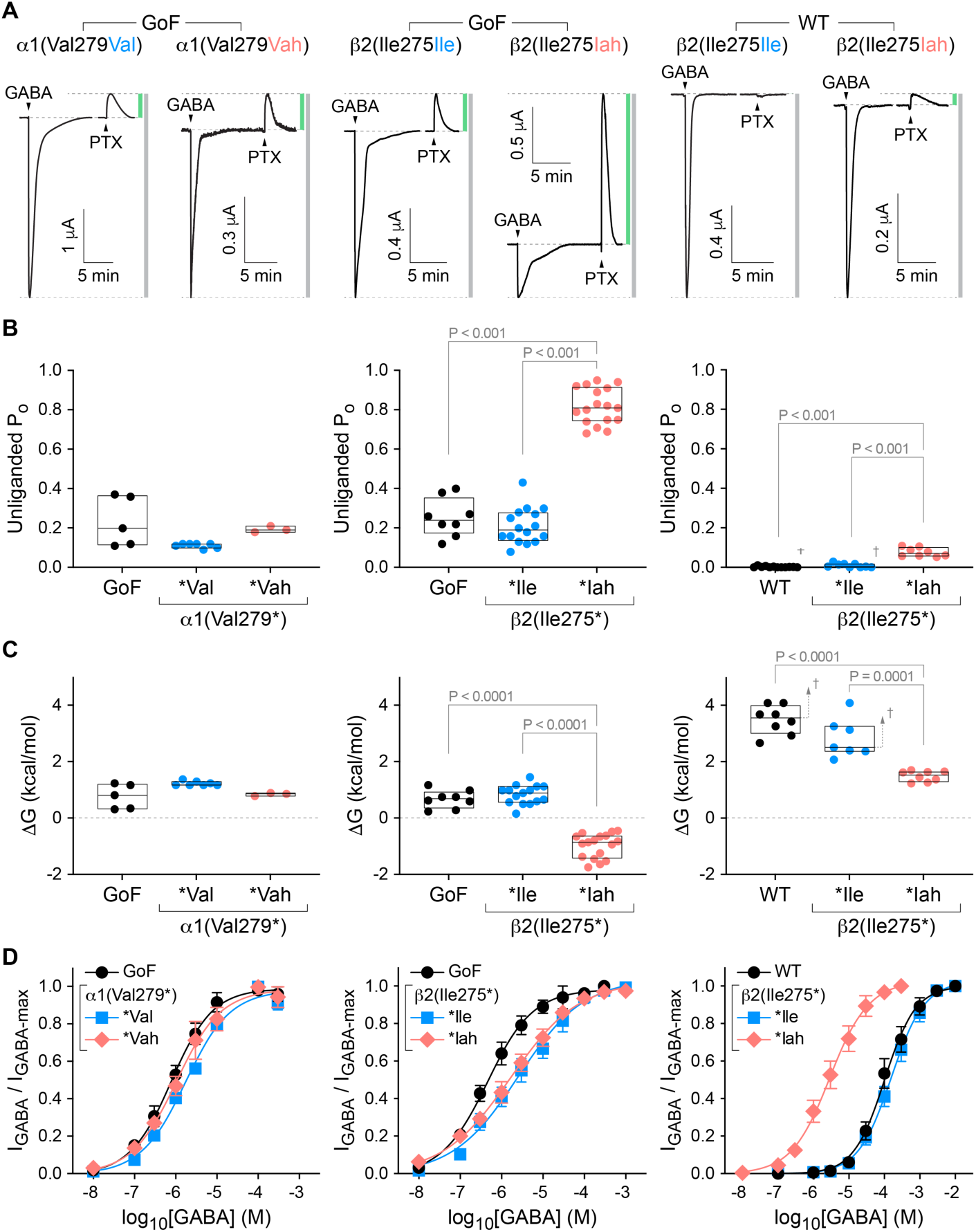
The β2 subunit main-chain Ile275:N-Pro273:O H-bond inhibits pore opening and is required to keep the unliganded channel closed. **(A)** Current responses to 10-20 s pulses (arrows) of saturating GABA or 1 mM PTX for α1β2γ2 (WT) or α1(Leu9′Thr)β2ψ2 (GoF) receptors after nonsense suppression incorporation of either the wild-type amino acid (Val, Ile) or its cognate α-hydroxy acid (Vah, Iah) at α1(Val279*) or β2(Ile275*). Vertical bars indicate the relative magnitude of PTX-sensitive (green) to total (gray) currents. **(B)** Unliganded open probability (P_o_) per oocyte estimated as the fraction of PTX-sensitive to total current. Box plots show median and interquartile intervals. P-values < 0.05 for Brown-Forsythe ANOVA with posthoc Dunnett’s T3 test shown. **(C)** Free energy difference between closed and open states in the absence of ligand for each oocyte computed from the estimated open probabilities in panel b. Box plots and P-values as in panel B. **(D)** Normalized concentration-response relations for GABA-elicited currents. Data are mean ± SEM across oocytes. Curves are the Hill equation fit to the means (Equation 2). See Tables S1-S2 for summary statistics, fit parameters and number of oocytes. ^†^For WT receptors, the reported unliganded open probabilities and free energies are upper and lower bounds, respectively, due to the limited ability to resolve small currents on the order of the noise.

### H-bond in β2 inhibits pore opening

We initially expected rather small functional effects from ablating individual H-bonds. Thus, we first evaluated these perturbations in the α1(Leu9′Thr)β2γ2 gain-of-function (GoF) background. The substitution α1(Leu9′Thr) in the main pore gate stabilizes the open state such that the channel open spontaneously in the absence of agonist with appreciable probability ^38^. This behavior allows relatively small changes in the closed-open equilibrium to be readily observed as changes in ionic current ^36,37^. Due to limited desensitization, the peak response of GoF receptors to saturating GABA is predicted to be close to maximal (i.e., an open probability of 1). Application of the pore blocker PTX blocks any spontaneously open channels and reveals the zero-current baseline (i.e., an open probability of 0). Thus, we estimated channel open probability by normalizing currents between the extreme levels elicited separately with PTX and saturating GABA (*I_Total_*) (**Figures 2C**, **3A**). Basal unliganded open probability (*P_o_*) is thus given by the ratio of PTX-sensitive to total current amplitude (*I_PTX_*/*I_Total_*), from which the closed-open free energy difference is computed as

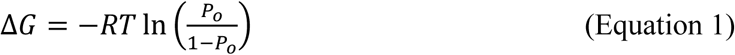

where *R* is the gas constant and *T* is temperature.

Nonsense suppression reintroduction of WT amino acids at TAG sites, α1(Val279Val) or β2(Ile275Ile), recapitulated GoF behavior including both basal *P*_*o*_ (**Figure 3A-C left, middle**) and GABA sensitivity (**Figure 3D left, middle**). Although there was a slight shallowing of the GABA concentration-response curves (CRCs) for nonsense suppression incorporation of residues at β2(Ile275*) as compared to GoF, this effect was minor in comparison to the effects we will focus on. Similarly, ablation of the predicted H-bond α1(Val279:N-Pro277:O) with the substitution α1(Val279Vah) had no effect on GoF behavior, suggesting that either this H-bond does not exist, or it is irrelevant to channel function. Given the location of the α1 subunit M2-M3 linker below the classical benzodiazepine site in the ECD and our previous observation that α1(Val279Ala) enhances diazepam modulation ^36^, we asked whether the α1(Val279:N-Pro277:O) H-bond was involved in allosteric modulation by benzodiazepines such as diazepam. Consistent with its lack of effect on GABA-elicited currents, α1(Val279Vah) also had no effect on allosteric modulation by diazepam (**Figure S3**).

In contrast, ablation of the predicted H-bond β2(Ile275:N-Pro273:O) with the substitution β2(Ile275Iah) massively increased the estimated basal *P*_*o*_as compared to GoF (**Figure 3A-B middle**), suggesting that this H-bond naturally inhibits channel opening by ∼1.8 kcal/mol (**Figure 3C middle**). We did not observe an associated increase in GABA sensitivity (i.e., left-shift of the CRC) (**Figure 3D middle**). However, we have previously observed that combinations of GoF and other sensitizing mutations can have non-additive effects on GABA sensitivity ^37^, possibly due to overlapping mechanisms for their sensitizing effects.

### H-bond in β2 required to stay closed

Based on the large effect of β2(Ile275Iah) in the GoF background, we examined substitutions at β2(Ile275*) in the α1β2γ2 WT background. First, β2(Ile275Ile) recapitulated WT behavior including little to no PTX-sensitive basal current (**Figure 3A-C right**) and normal GABA sensitivity (**Figure 3D right**). Consistent with our observation in the GoF background, ablation of the predicted H-bond β2(Ile275:N-Pro273:O) in a WT background with the substitution β2(Ile275Iah) increased the basal *P*_*o*_to the extent that channels were spontaneously open ∼7% of the time (**Figure 3A-B right**). Although we cannot rule out an effect on desensitization, the observation of spontaneous unliganded opening in a WT background shows that elimination of the β2(Ile275:N-Pro273:O) H-bond strongly promotes channel opening. For WT or β2(Ile275Ile), PTX-sensitive currents were sufficiently small to be either unmeasurable or difficult to disambiguate from noise. Thus, our estimated basal *P*_*o*_ of ∼0.002 for these channels represents an upper limit, and conversely the calculated Δ*G* values reflect a lower limit. Nonetheless, we estimate that the two H-bonds β2(Ile275:N-Pro273:O) inhibit WT channel opening by at least ∼1.5 kcal/mol (**Figure 3C right**), similar to our estimation for GoF channels. Furthermore, this H-bond stabilizes a non-conducting conformation to the extent that it is required to keep the unliganded channel closed with high probability.

### H-bond in β2 is state-dependent

In the WT background, elimination of the H-bond β2(Ile275:N-Pro273:O) with the substitution β2(Ile275Iah) enhanced apparent GABA sensitivity by left-shifting GABA CRCs ∼50-fold (**Figure 3D right**). This suggests that breaking of this H-bond enhances channel opening by the same or a similar mechanism as normal activation by GABA, in that the expected coupling between the pore gate in the TMD and the agonist sites in the ECD is qualitatively preserved (i.e., agonists have higher affinity for open versus closed states). Thus, we hypothesize that normal channel activation by GABA involves breaking of the main-chain H-bond β2(Ile275:N-Pro273:O).

To further explore this idea, we examined the spatial proximity of the donor and acceptor atoms as a proxy for H-bond likelihood during all-atom molecular dynamics simulations of α1β2γ2 receptors. Comparison of simulations for antagonist-bound (i.e., closed) and GABA-bound (i.e., activated/desensitized) conformations indicates that channel activation is associated with an increased donor-acceptor separation for β2(Ile275:N-Pro273:O), consistent with a reduced H-bond likelihood in open versus closed states (**Figure 4A-B**). Strikingly, donor-acceptor distance distributions for the analogous H-bonds in α1 and γ2 subunits were relatively independent of ligation state and largely similar to that for β2 subunits in a closed state. If anything, H-bond distances in α1 and γ2 subunits were slightly shortened on average upon receptor activation by GABA. These results are consistent with our observation that the H-bond α1(Val279:N-Pro277:O) is not involved in channel function. Taken together, these observations strongly suggest that specifically the main-chain H-bond β2(Ile275:N-Pro273:O) stabilizes a closed conformation and that this bond is broken during channel opening (**Figure 4C**).

**Figure 4.**
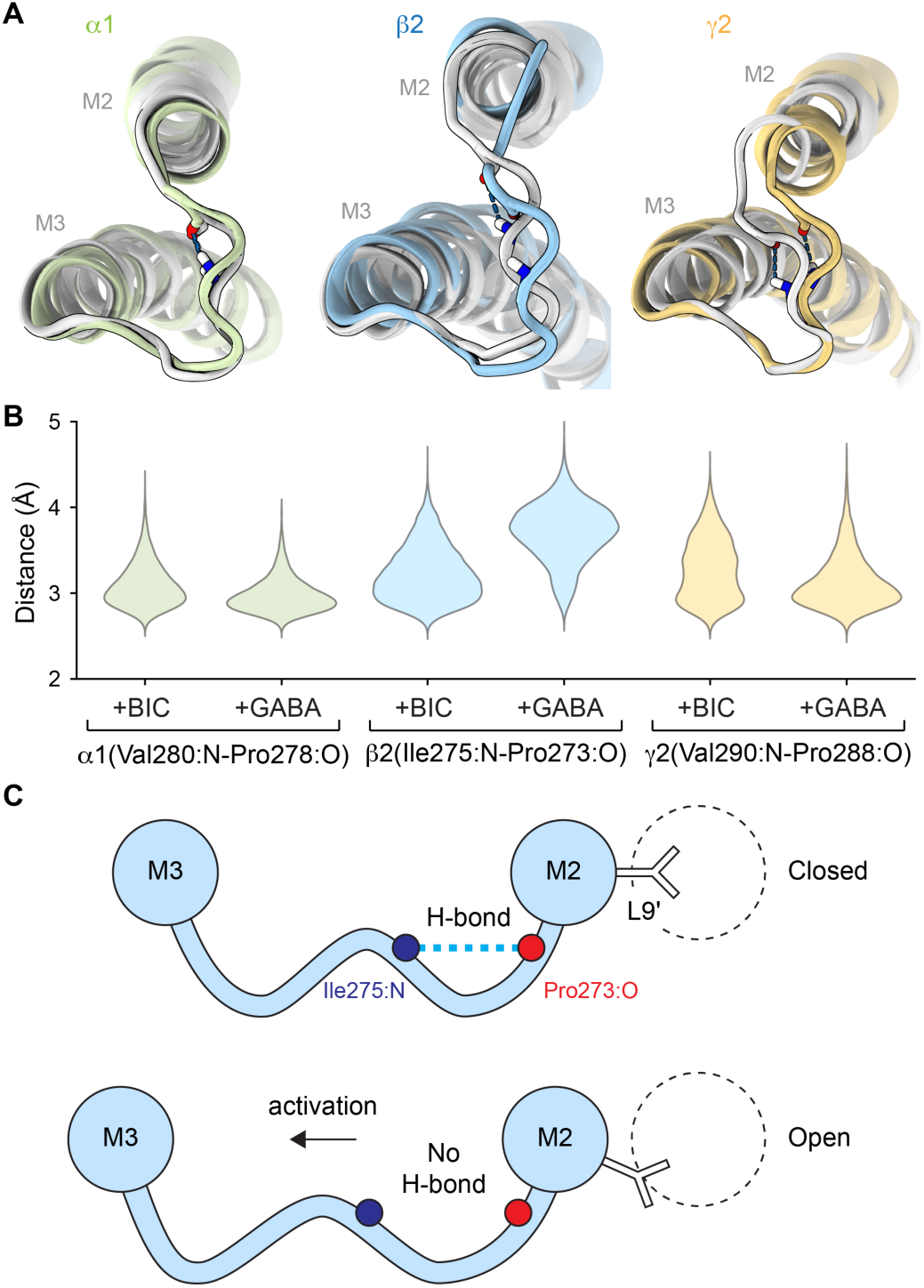
A state-dependent main-chain H-bond in the β2 subunit M2-M3 linker is a hair-trigger for channel opening. **(A)** Representative poses for M2-M3 linkers from MD simulations of antagonist-bound (i.e., closed; PDB 6X3S; white) and GABA-bound (i.e., activated; PDB 6X3Z; color) human GABA_A_R complexes, aligned per subunit on local M2 and M3 helices. H-bonds α1(Val280:N-Pro278:O), β2(Ile275:N-Pro273:O) and ψ2(Val290:N-Pro288:O) are indicated with dashed lines, and participating atoms are shown as sticks. **(B)** Distance distributions between donor and acceptor atoms for the H-bonds depicted in panel a from MD simulations of either antagonist-bound (+BIC, bicuculine) or GABA-bound (+GABA) complexes. Distributions are aggregate across four replicates of 500 ns simulations (Figure S5). Note, human α1(Val280:N-Pro278:O) is analogous with rat α1(Val279:N-Pro277:O) (Figure S1). **(C)** Cartoon illustrating the proposed breaking of the state-dependent H-bond β2(Ile275:N-Pro273:O) within the M2-M3 linker during channel opening.

For WT receptors, the maximal *P*_*o*_ upon activation by saturating GABA is ∼0.8 ^39^. Given a basal unliganded *P*_*o*_ of ∼0.002 (**Figure 3B right**) ^40^, the chemical energy from GABA binding results in a shift of the closed-open free energy difference by ∼4.5 kcal/mol (see Eq. 1). Thus, breaking the main-chain H-bond β2(Ile275:N-Pro273:O) in each of two subunits accounts for ∼1/3^rd^ of the channels total activation energy. Note that this is a lower limit in the case that we have overestimated the basal *P_o_*.

## Discussion

Here we show that a main-chain H-bond within the β2 M2-M3 linker inhibits pore opening, is required to keep the unliganded channel closed at rest, and that breaking of this bond during activation by GABA accounts for ∼1/3^rd^ of the energy derived from GABA binding. Structures suggest that this H-bond stabilizes a crimp or shallow hairpin in the M2-M3 linker backbone predicted to limit linker flexibility (**Figure 1B**). We propose that increased flexibility of the β2 M2-M3 linker upon breaking this bond contributes to the physical basis for channel activation. Consistent with this idea, the substitution β2(Ile275Ala), which we predict to also increase linker flexibility by removing side-chain volume and reducing hydrophobicity near the center of the linker, has a similar functional signature: enhanced opening with a constitutive unliganded PTX-sensitive current and increased apparent GABA sensitivity ^37^.

Ablation of the analogous main-chain H-bond within the α1 M2-M3 linker had no effect on channel function. Strikingly, this also qualitatively parallels prior observations of asymmetric effects for alanine substitutions predicted to increase M2-M3 linker flexibility in α1, β2 or γ2 subunits ^37^. Namely, β2(Ile275Iah) or β2(Ile275Ala) promote channel opening and prevent the channel from remaining closed, whereas the analogous mutations α1(Val279Vah), α1(Val279Ala), or γ2(Val290Ala) either have no effect or inhibit channel gating. Although other studies have observed subunit-specific effects of M2-M3 linker mutations ^31,33^, a coherent picture for the physical basis underlying this asymmetry is unclear. We hypothesize that linker flexibility is differentially transduced to channel gating at distinct subunits or interfaces. Asymmetry in the flexibility of M2 helices was also observed for cysteine cross-linking which indicates that the pore-lining M2 helices in β subunits are more radially mobile than those in α subunits ^41,42^. Consistent with these observations, comparison of antagonist-and GABA-bound cryo-EM structures show that activation involves relatively larger radial motions of M2-M3 linkers in β2 subunits as compared to α1 or γ2 subunits ^23,34,35^. Taken together, these results strongly suggest that flexibility of the β2 subunit M2-M3 linkers has a special role in channel activation as compared to the linkers in α1 or γ2 subunits, possibly due to their physical location at the GABA-binding inter-subunit interfaces.

Static structures of synaptic GABA_A_Rs enable rough prediction of the main-chain H-bonds investigated here. However, not only do they not make clear their mechanistic importance, they also do not completely clarify their existence. Based on default distance and rotamer cutoffs in programs such as ChimeraX ^43^, the investigated H-bonds are identified in some distinct subunits within some structures, but not all subunits/structures. It is unclear to what extent this reflects a lack of resolution in more mobile loop regions and/or dynamics of these bonds. MD simulations indicate a preference for donor-acceptor distances of just below three Angstroms (except for β2 subunits with GABA bound) (**Figure 4A**), typical for N·O H-bonds in proteins ^44,45^. However, non-static donor-acceptor distance distributions exhibit tails extending to over four Angstroms, consistent with a dynamic nature to the electrostatic strength of these main-chain H-bonds which may obfuscate their identification in static structures.

In conclusion, a main-chain H-bond within the β2 subunit M2-M3 linker is a critical component of the channel’s gating mechanism. We predict that this bond inhibits pore opening by limiting M2-M3 linker flexibility, and that breaking of this bond, or its conversion from stronger to weaker average strength, accounts for ∼1/3^rd^ of the activation energy supplied upon binding of GABA at both agonist sites. Other main-chain interactions in gating loops may also be important aspects of the physical basis for the behavior of GABA_A_Rs as well as other pLGICs. However, their existence and/or contribution to channel function is typically not obvious from static structures and remains to be determined. Lastly, it is notable that this hair-trigger mechanism for rapid channel opening involves only main-chain atoms, and thus is largely immune to genetic variation.

## Methods

### Animals

Mature female *Xenopus laevis* frogs were obtained from Nasco and housed in the University of Texas animal facility. Frog care and surgery followed the ARRIVE guidelines and an IACUC-approved protocol. Frogs were housed in tanks at a density of no more than 2 frogs per gallon. The water in the tanks is continuously drained and passed through a series of chemical, biological and UV filters before being returned. *Xenopus laevis* oocytes were harvested from frogs under tricaine anesthesia. A piece of ovary was removed from the frog, and placed in isolation media (108 mM NaCl, 2 mM KCl, 1 mM EDTA, 10 mM HEPES, pH = 7.5).

### Mutagenesis and in vitro transcription

DNA for wild-type and mutant GABA_A_R rat α1, β2, and γ2 subunits was subcloned in the pUNIV vector ^46^. The mature protein numeration for β2 and γ2 subunits is the same for rat and human, but the numeration for the rat α1 subunit is (human numeration -1) for most of the subunit **(Figure S1**). For ncAA incorporation, the codon of the residue of interest was replaced by the TAG stop codon (in contrast to TGA stop codons for each subunit). Mutations were introduced using QuikChange II (Qiagen) or by GenScript and confirmed by sequencing of the entire subunit. Sequences for all constructs are provided in supplementary information. Complementary RNA (cRNA) for each construct was generated (mMessage mMachine T7, Ambion), quantified (Qubit, ThermoFisher Scientific) and quality assessed (TapeStation, Agilent) prior to injection in *Xenopus laevis* oocytes.

### ncAA synthesis

For nonsense suppression in GABA_A_ subunits, we used TAG mutants of the α1 and β2 GABA_A_ subunits and PylT tRNAs in *Xenopus laevis* oocytes. PylT lacking the two terminal CA nucleotides was synthetized by Integrated DNA Technologies, Inc., folded and misacylated as previously described ^6^. Leu-, Val-, α-hydroxy Leu-(Lah), and α-hydroxy Val-(Vah) pdCpA-substrates were synthesized according to published procedures ^6^.

### Isolation and injection of *Xenopus laevis* oocytes

A portion of the ovary was extracted from the frog, after which oocytes were manually isolated from the thecal and epithelial layers using forceps and then incubated in a collagenase buffer (0.5 mg/mL collagenase from *Clostridium histolytic,* 83 mM NaCl, 2 mM KCl, 1 mM MgCl_2_, 5 mM HEPES) to remove the follicular layer. Oocytes were injected with 12 ng of total cRNA for α1, β2, and γ2 subunits (wild-type or mutants) in a 1:1:10 ratio ^47^ (Nanoject, Drummond Scientific). When injecting the TAG mutant of a subunit, 125 ng of tRNA was mixed with the cRNA encoding the GABA_A_ subunits. Oocytes were incubated in a sterile incubation solution (88 mM NaCl, 1mM KCl, 2.4 mM NaHCO_3_, 19 mM HEPES, 0.82 mM MgSO_4_, 0.33 mM Ca(NO_3_)_2_, 0.91 mM CaCl_2_, 10,000 units/L penicillin, 50 mg/L gentamicin, 90 mg/L theophylline, and 220 mg/L sodium pyruvate, pH=7.5) at 16 °C.

### TEVC recording in oocytes

Currents from expressed channels 1-3 days post-injection were recorded in two-electrode voltage clamp (Oocyte Clamp OC-725C, Warner Instruments) and digitized using a PowerLab 4/30 system (ADInstruments). Oocytes were held at -70 mV and perfused continuously (2 mL/min) with ND96 buffer (96 mM NaCl, 2 mM KCl, 1 mM CaCl_2_, 1 mM MgCl_2_, 5 mM HEPES, pH 7.5) or ND96 buffer containing picrotoxin (PTX), GABA, or diazepam (DZ). PTX was diluted from a 0.5 M stock solution in DMSO. DZ was diluted from a 10 mM stock solution in DMSO. GABA was diluted from 300 mM stock solution in water. The recording protocol was as follows: a 10 s pulse of PTX was followed by a series of 20-40 s pulses of increasing concentrations of either GABA or DZ and a final 10 s pulse of PTX. Pulses were sufficiently long to resolve the peak response and inter-pulse intervals were 5–15 min to allow washout with buffer and currents to return to baseline. A representative trace is shown in **Figure 2C**. Current traces were analyzed with custom scripts in MATLAB (Mathworks). GABA or DZ concentration-response curves (CRCs) were fit with the Hill equation:

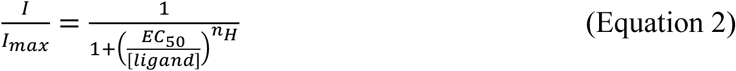

where *I* is the magnitude of the ligand-elicited current, [*ligand*] is GABA or DZ concentration, *EC*_50_ is the concentration eliciting a half-maximal response, and *n_H_* is the Hill slope.

For α1(Leu9’Thr)-containing receptors, we often observed that the spontaneous unliganded current decreased in magnitude with time in a nonlinear fashion, decreasing more rapidly at the beginning of the recording and reaching a nearly stable baseline in the latter period of the recording. This is unlikely to be accounted for by changes in leak current alone as PTX consistently reduced the current to a similar level at the beginning and end of the recording. Thus, we assume that this rundown of the spontaneous current reflects a reduction in the pool of active channels at the membrane. Therefore, for α1(Leu9’Thr)-containing receptors, we first fit a spline to the spontaneous current baseline and subtracted this fit from the raw current. To normalize out the time-dependent loss of active channels, we divided the baselined currents by the magnitude of the spline approximation of the spontaneous current baseline (i.e., our proxy for the number of active channels). Finally, we normalized the resulting detrended currents between the final PTX-and maximal GABA-elicited current levels as an approximation for open probability. See **Figure S2** for a depiction of this protocol which reasonably accounts for most of the observed rundown as evidenced by the very similar magnitude PTX-elicited responses at the beginning and end of the detrended recording despite not enforcing this a priori. Most importantly, this detrending has almost no effect on the relative magnitude of the final responses to saturating GABA and PTX at the end of the recording where the baseline is largely stable, and thus does not affect our measure of unliganded open probability. The only effect this procedure has is to reduce the relative magnitude of current responses to low GABA concentrations towards the beginning of the recording, effectively reducing the foot of the concentration-response curves in a way that we believe better reflects the actual GABA sensitivity of these channels. Regardless, this adjustment is irrelevant for all our major conclusions.

### Molecular dynamics simulations

MD simulations were as described previously ^27^. Briefly, each structure of the α1β2ψ2 GABA_A_ receptor in the presence of relevant ligands was placed in an MD simulation box with dimensions 127 × 127 × 163 Å^3^, embedded in a bilayer of 400 1-palmitoyl-2-oleoyl-sn-glycero-3-phosphocholine (POPC) molecules, and subsequently solvated with TIP3P water ^48^ and 150 mM NaCl in CHARMM-GUI ^49^. The CHARMM36m forcefield ^50^ was used to describe the protein. Ligand parameters were generated with CGenFF in CHARMM-GUI ^51^. Simulations were performed using GROMACS 2019.3 ^52^. The temperature was kept at 300 K using a velocity-rescaling thermostat ^53^. Parrinello-Rahman pressure coupling ^54^ ensured constant pressure, the particle mesh Ewald algorithm ^55^ was used for long-range electrostatic interactions, and hydrogen-bond lengths were constrained using the LINCS algorithm ^56^. After each system was energy-minimized, sequential 10 ns equilibration steps were performed with gradual release of position restraints on heavy, backbone, and Cα atoms. The bicuculline ligands were restrained during equilibration. Four replicates of ∼500 ns unrestrained simulations were then generated, and frames were analyzed every 2 ns, for a total of 1000 samples (4 replicates × 250 frames) for each system. Local RMSDs for M2-M3 linkers indicate convergence of simulations in the region of interest (**Figure S4**). Distances between hydrogen-bonding atoms were calculated using MDAnalysis ^57^.

Simulations in the presence of bicuculline or GABA were initiated from corresponding cryo-EM structures (PDB IDs: 6X3S or 6X3Z). Alternative analyses of the GABA-bound simulations were also reported previously ^27^. Simulations under both conditions (bicuculline and GABA) are available on Zenodo ^58^.

### Statistical analysis

Data was obtained from oocytes from at least two different frogs for each experimental group. Summary data was analyzed using Prism 10 (GraphPad). Symbols and error bars are mean ± SEM, and box plots show median and interquartile intervals. Further details can be found in the figure legends and Tables S1-S2. Where applicable, we applied One-way Brown-Forsythe ANOVA (as the standard deviations were not equal among groups) followed by Dunnett’s T3 multiple comparisons test. Irrespective of P-values, we focus only on relatively large effects.

## Acknowledgements

This research was supported by NIH grants R01GM148591 to M.P.G-O. and R35GM148239 to C.A.A., and by Swedish Research Council (VR) grants 2019-02433 and 2021-05806 to R.J.H. and E.L. Additional support was provided by the Sven & Lily Lawski Foundation to S.E.L., and by the Knut & Alice Wallenberg Foundation and Swedish e-Science Research Centre to R.J.H. and E.L.

## Author contributions

M.P.G.-O. conceived and supervised this work. J.D.G. carried out, and C.A.A supervised, the ncAA synthesis. C.M.B, M.P.G.-O. and C.A.A. established ncAA protocols for TEVC recordings.

C.M.B and N.G.D. carried out, and C.M.B analyzed, the TEVC recordings. S.E.L and Y.Z. carried out and analyzed, and R.J.H and E.L. supervised, the molecular dynamics simulations. C.M.B, M.P.G.-O. and S.E.L. produced the visualizations. C.M.B and M.P.G.-O. wrote the manuscript.

---

**Table S1.**
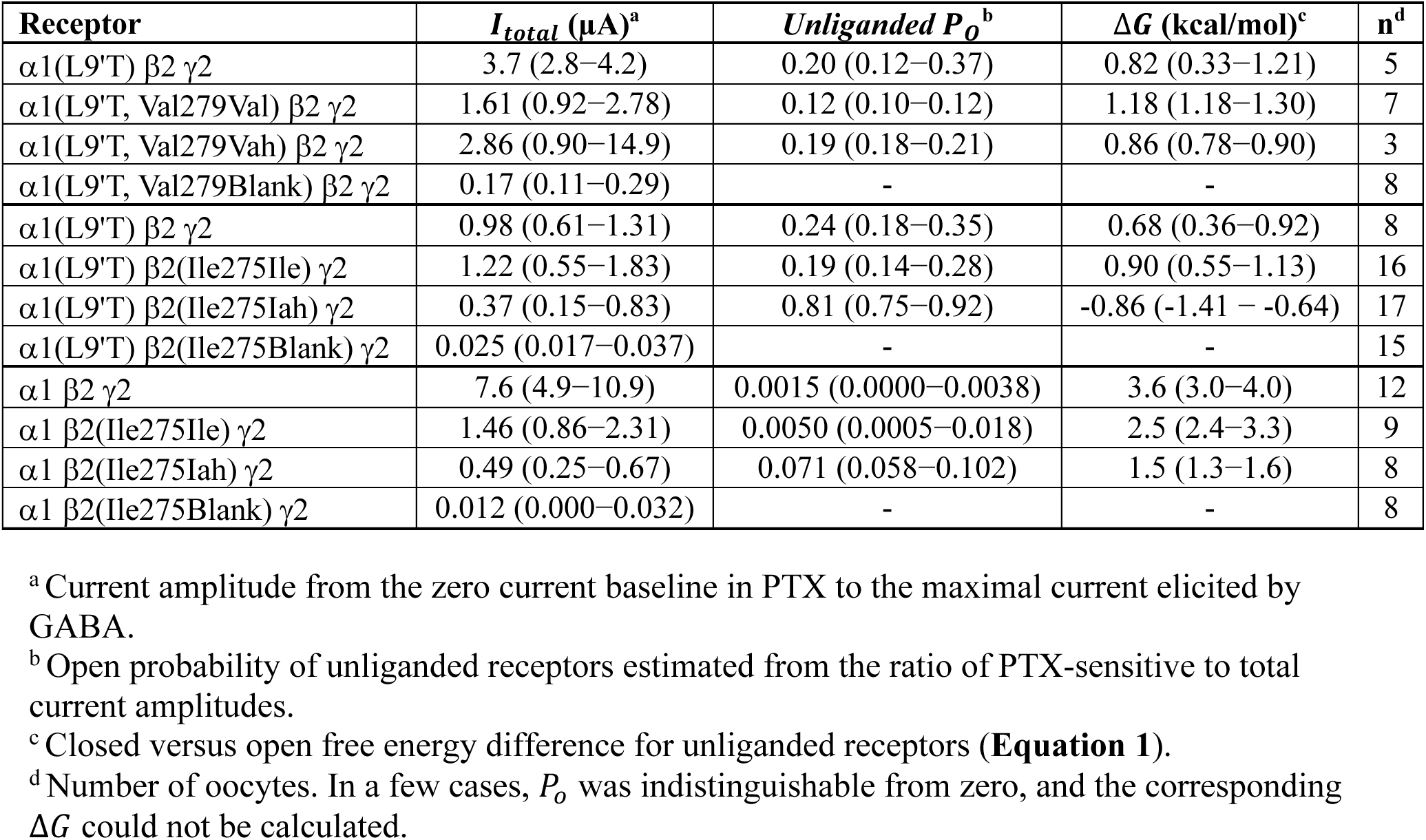
Summary of results from. **Figures 2-3**, **expressed as median (interquartile range)**

**Table S2.**
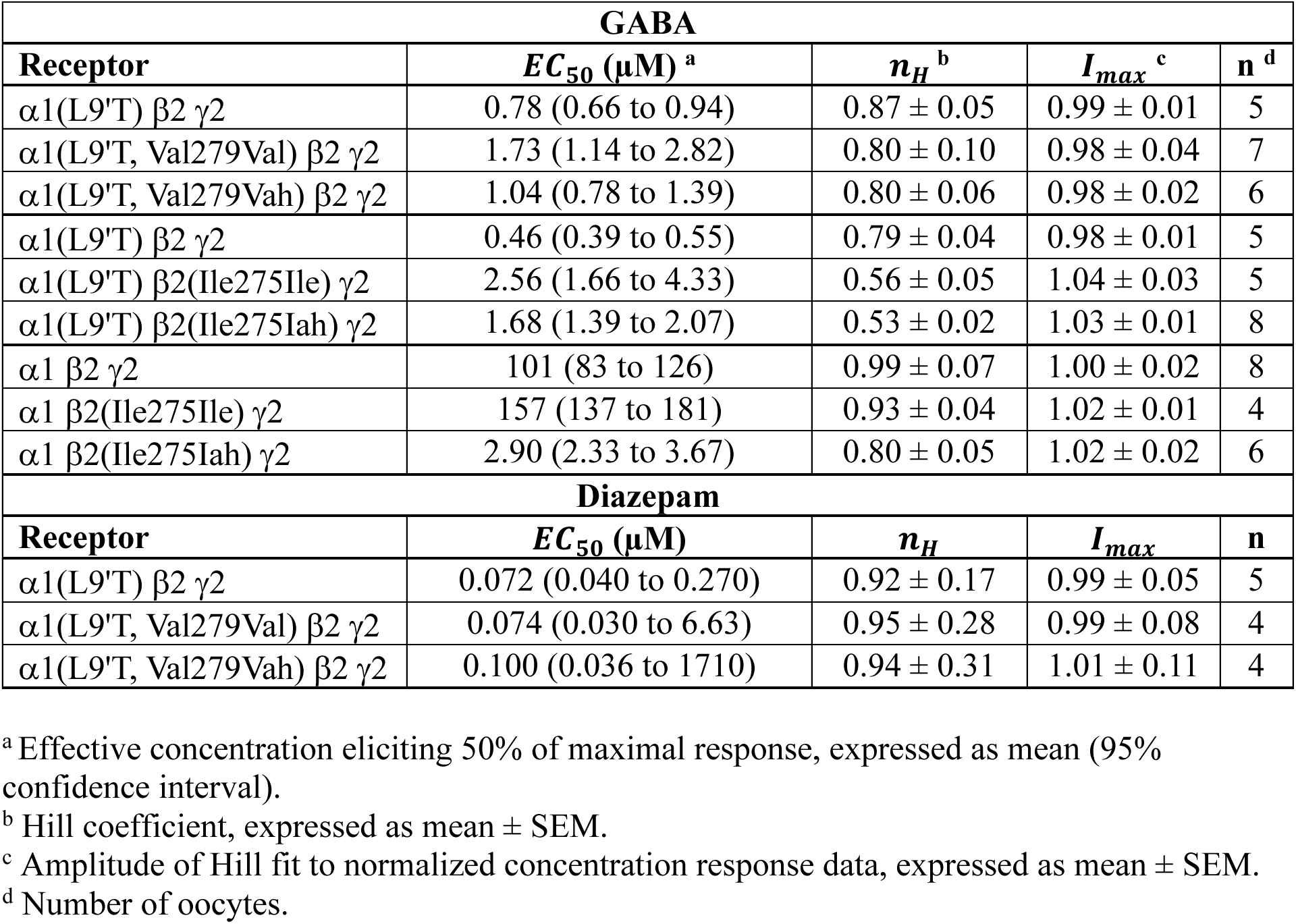
Parameters for Hill equation (Equation 2) fits to GABA and diazepam concentration-response data.

**Figure S1.**
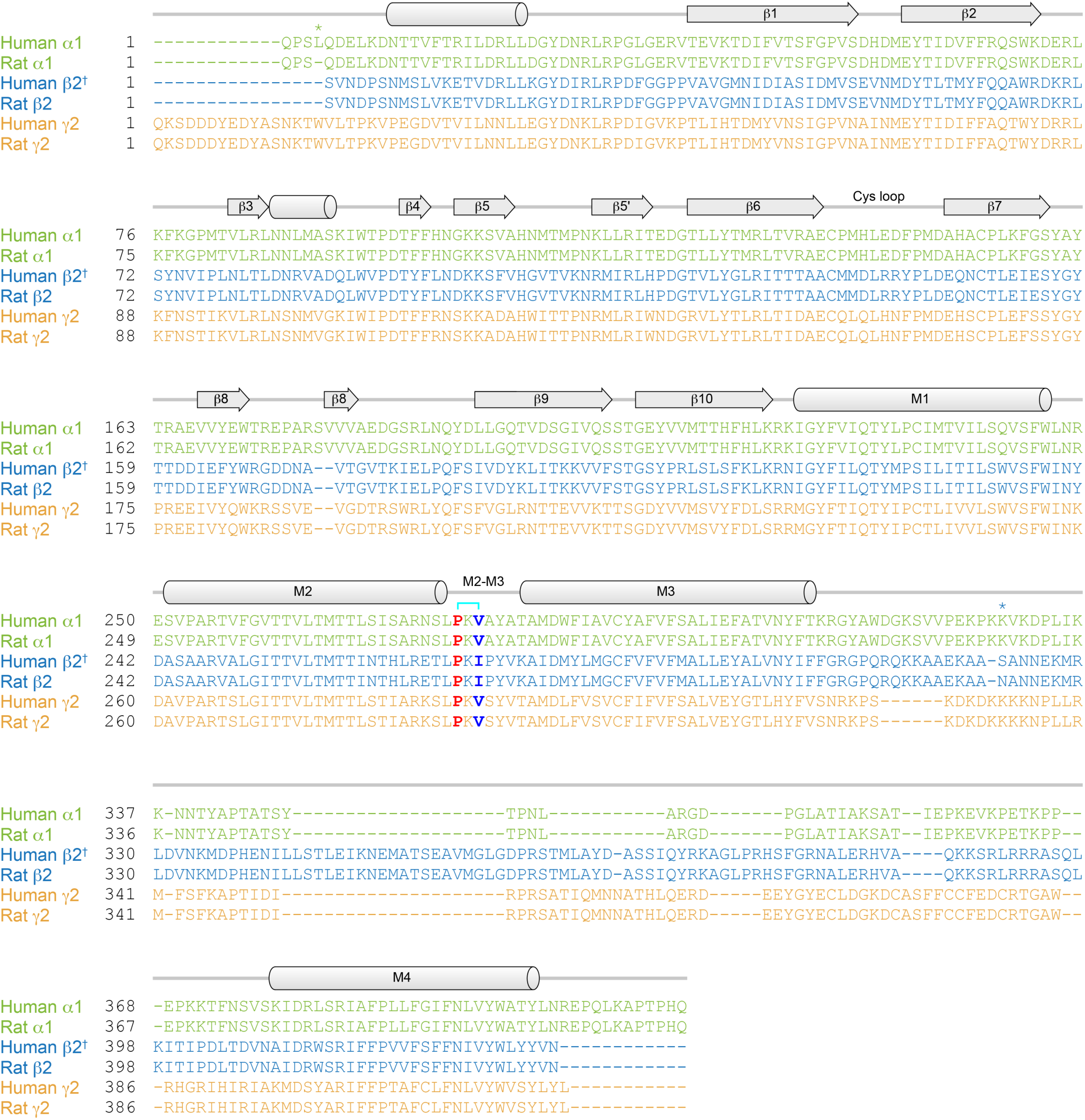
Human and rat sequences of GABA_A_ subunits. Human and rat sequences are either identical (ψ2) or differ by either a single residue gap in the N-terminus (α1) or a single residue substitution in the M3-M4 linker (β2). Residue numbering is for the mature protein. Notably, human and rat residue numbering is shifted by one for most of the α1 subunit. Approximate locations of helix and sheet secondary structures are indicated above the sequences. The main-chain H-bond donor (blue) and acceptor (red) residues in the M2-M3 linkers are bolded. Protein sequence identifiers from UniProt are P14867 (human α1), P62813 (rat α1), P47870-1 (human β2, ^†^short isoform), P63138 (rat β2), P18507 (human ψ2), and Q6PW52 (rat ψ2). The canonical β2 subunit (P47870-2, long isoform) includes an additional stretch of 38 amino acids in the intracellular M3-M4 linker compared to the short isoform.

**Figure S2.**
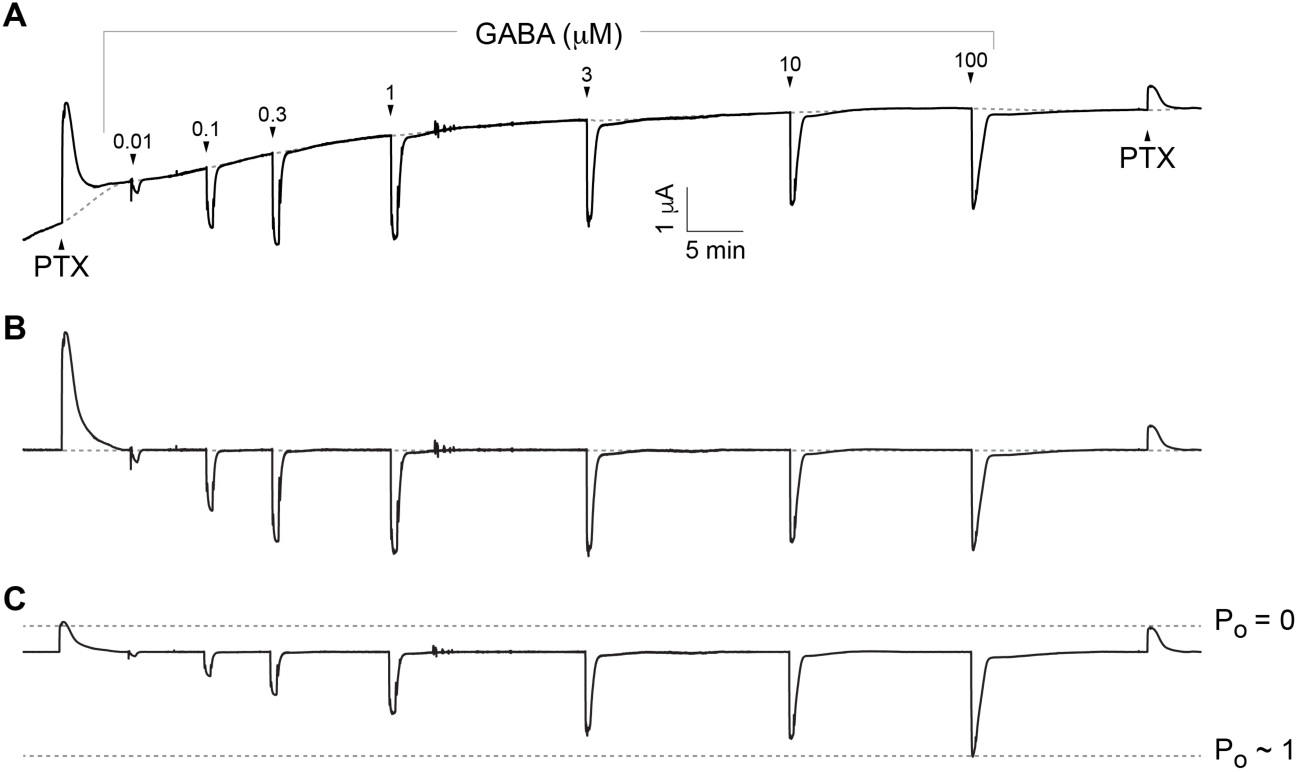
Detrending rundown of current time series traces as described in methods. **(A)** Current trace for α1(Leu9′Thr,Val279Vah)β2γ2 receptors illustrating pulse protocol. Arrows indicate approximate onset of pulses of either 1 mM PTX or increasing concentrations of GABA. Dashed gray line is a spline fit to the basal current level between ligand applications. Rundown of constitutive PTX-sensitive current amplitude interpreted as a time-dependent loss of active channels. **(B)** Baselined current after subtraction of the spline approximation to the baseline constitutive current from the raw current time series in panel A. **(C)** Detrended current after dividing the baselined current in panel B by the magnitude of the spline approximation to the constitutive current (i.e., a value proportional to the number of active channels) and finally rescaling to match the original amplitude of the final response to PTX. This procedure reasonably accounts for the observed rundown in the number of active channels as evidenced by the similar PTX-sensitive current amplitudes at the beginning and end of the detrended trace despite not enforcing this a priori. Most importantly, this detrending has almost no effect on the relative magnitude of the final responses to saturating GABA and PTX at the end of the recording from which our primary conclusions are drawn.

**Figure S3.**
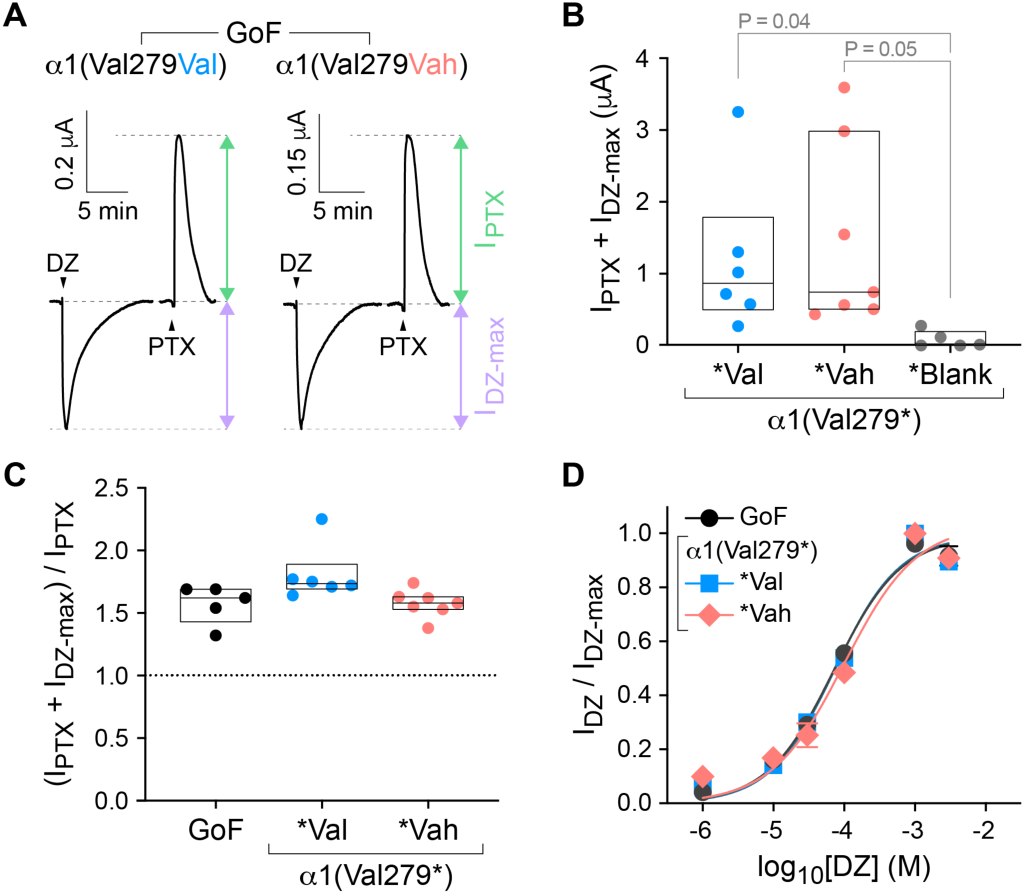
H-bond in α1 does not affect diazepam (DZ) modulation. **(A)** Current responses to 10-20 s pulses (arrows) of DZ eliciting a maximal response or 1 mM PTX for α1(Leu9′Thr)β2γ2 (GoF) receptors after nonsense suppression incorporation of either the wild-type amino acid (Val) or its cognate α-hydroxy acid (Vah) at α1(Val279*). Vertical bars indicate the relative magnitude of PTX-sensitive (green) to DZ-elicited (purple) currents. **(B)** per oocyte (left-to-right: n = 5, 7, 3). Box plots show median and interquartile intervals. P-values <= 0.05 for Brown-Forsythe ANOVA with posthoc Dunnett’s T3 test shown. **(C)** Maximal fold-potentiation of basal unliganded current by DZ (left-to-right: 8, 16, 17). **(D)** Normalized concentration-response relations for DZ-elicited currents. Data are mean ± SEM across oocytes. Curves are the Hill equation fit to the means (Equation 2). See Table S2 for fit parameters and number of oocytes.

**Figure S4.**
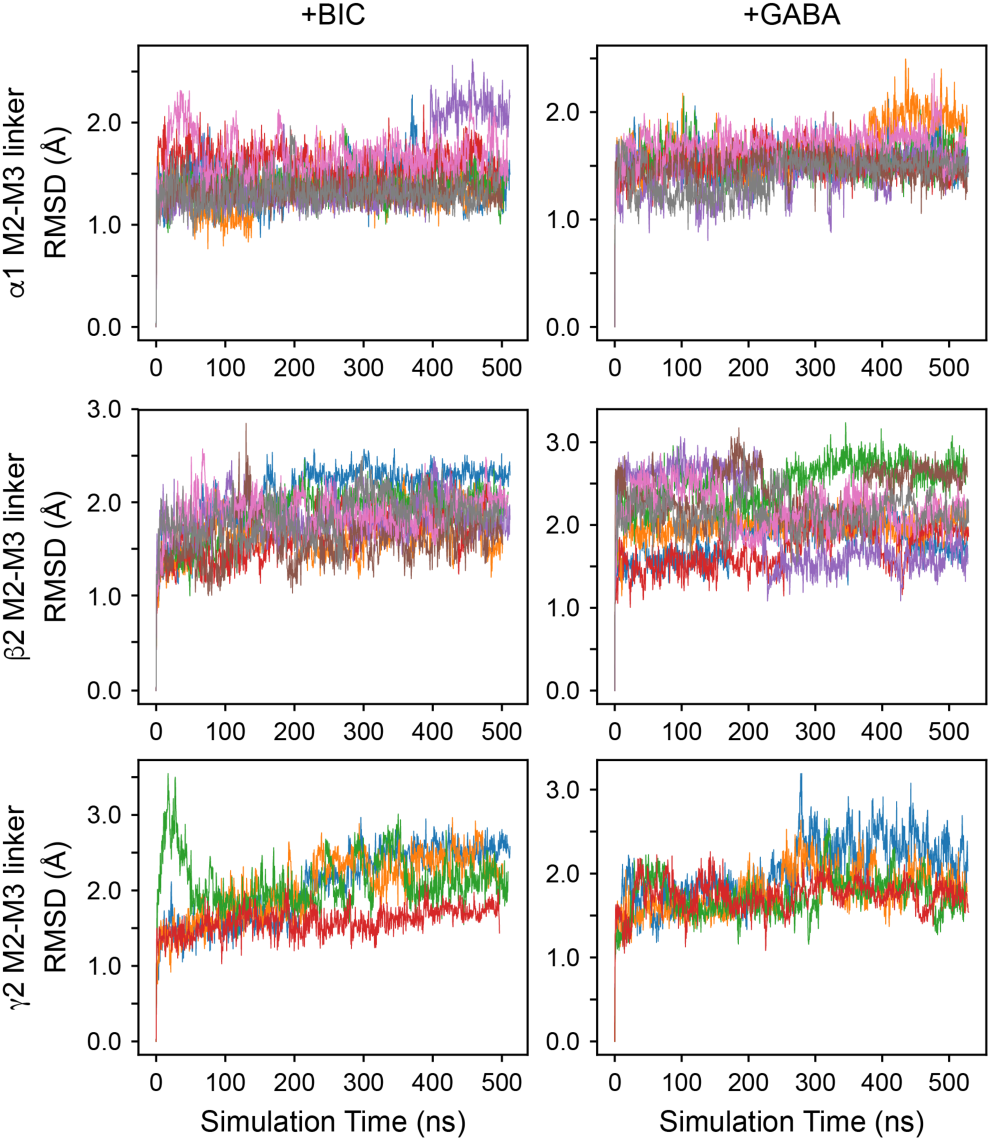
Convergence of MD simulations for M2-M3 linkers in each subunit. Left panels, antagonist-bound complexes (+BIC, bicuculine; PDB 6X3S). Right panels, GABA-bound (+GABA; PDB 6X3Z) complexes. Distinct chains for each of four replicates are colored differently.

**Figure S5.**
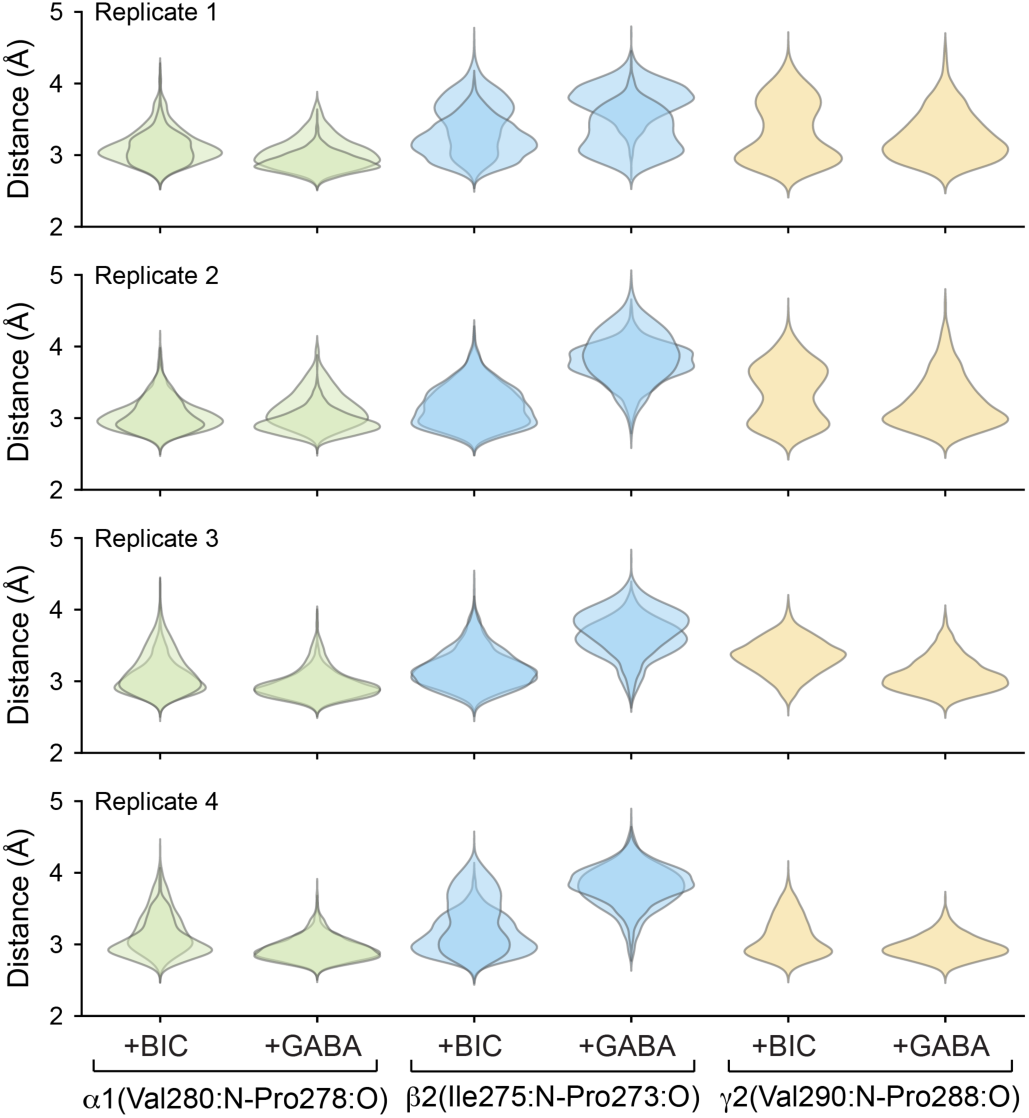
Chain-specific distance distributions for pairs of donor/acceptor atoms predicted to form a main-chain H-bond within the M2-M3 linker from MD simulations. Results shown are for either antagonist-bound (+BIC, bicuculine; PDB 6X3S) or GABA-bound (+GABA; PDB 6X3Z) complexes. The distributions are shown per replica, with overlapping distributions corresponding to the two distinct chains for α1 and β2 subunits within the receptor.

## References

1. Infield, D. T., Schene, M. E., Galpin, J. D. & Ahern, C. A. Genetic code expansion for mechanistic studies in ion channels: an (un)natural union of chemistry and biology. Chem. Rev. (2024) doi:10.1021/acs.chemrev.4c00306.

2. Van Arnam, E. B. & Dougherty, D. A. Functional probes of drug-receptor interactions implicated by structural studies: Cys-loop receptors provide a fertile testing ground. J. Med. Chem. 57, 6289–6300 (2014).

3. Sereikaitė, V. et al. Probing Backbone Hydrogen Bonds in Proteins by Amide-to-Ester Mutations. Chembiochem 19, 2136–2145 (2018).

4. Leisle, L. et al. Backbone amides are determinants of Cl-selectivity in CLC ion channels. Nat. Commun. 13, 7508 (2022).

5. Lynagh, T. et al. A selectivity filter at the intracellular end of the acid-sensing ion channel pore. eLife 6, (2017).

6. Infield, D. T. et al. Main-chain mutagenesis reveals intrahelical coupling in an ion channel voltage-sensor. Nat. Commun. 9, 5055 (2018).

7. Nagaoka, Y., Shang, L., Banerjee, A., Bayley, H. & Tucker, S. J. Peptide backbone mutagenesis of putative gating hinges in a potassium ion channel. Chembiochem 9, 1725– 1728 (2008).

8. Lu, T. et al. Probing ion permeation and gating in a K+ channel with backbone mutations in the selectivity filter. Nat. Neurosci. 4, 239–246 (2001).

9. Blum, A. P., Gleitsman, K. R., Lester, H. A. & Dougherty, D. A. Evidence for an extended hydrogen bond network in the binding site of the nicotinic receptor: role of the vicinal disulfide of the alpha1 subunit. J. Biol. Chem. 286, 32251–32258 (2011).

10. Blum, A. P., Lester, H. A. & Dougherty, D. A. Nicotinic pharmacophore: the pyridine N of nicotine and carbonyl of acetylcholine hydrogen bond across a subunit interface to a backbone NH. Proc Natl Acad Sci USA 107, 13206–13211 (2010).

11. Cashin, A. L., Petersson, E. J., Lester, H. A. & Dougherty, D. A. Using physical chemistry to differentiate nicotinic from cholinergic agonists at the nicotinic acetylcholine receptor. J. Am. Chem. Soc. 127, 350–356 (2005).

12. Post, M. R., Lester, H. A. & Dougherty, D. A. Probing for and Quantifying Agonist Hydrogen Bonds in α6β2 Nicotinic Acetylcholine Receptors. Biochemistry 56, 1836– 1840 (2017).

13. England, P. M., Zhang, Y., Dougherty, D. A. & Lester, H. A. Backbone mutations in transmembrane domains of a ligand-gated ion channel: implications for the mechanism of gating. Cell 96, 89–98 (1999).

14. Gleitsman, K. R., Kedrowski, S. M. A., Lester, H. A. & Dougherty, D. A. An intersubunit hydrogen bond in the nicotinic acetylcholine receptor that contributes to channel gating. J. Biol. Chem. 283, 35638–35643 (2008).

15. Gleitsman, K. R., Lester, H. A. & Dougherty, D. A. Probing the role of backbone hydrogen bonding in a critical beta sheet of the extracellular domain of a cys-loop receptor. Chembiochem 10, 1385–1391 (2009).

16. Rienzo, M., Lummis, S. C. R. & Dougherty, D. A. Structural requirements in the transmembrane domain of GLIC revealed by incorporation of noncanonical histidine analogs. Chem. Biol. 21, 1700–1706 (2014).

17. Pless, S. A. & Sivilotti, L. G. A tale of ligands big and small: an update on how pentameric ligand-gated ion channels interact with agonists and proteins. Curr. Opin. Physiol. 2, 19–26 (2019).

18. Olsen, R. W. & Sieghart, W. International Union of Pharmacology. LXX. Subtypes of gamma-aminobutyric acid(A) receptors: classification on the basis of subunit composition, pharmacology, and function. Update. Pharmacol. Rev. 60, 243–260 (2008).

19. Sente, A. et al. Differential assembly diversifies GABAA receptor structures and signalling. Nature 604, 190–194 (2022).

20. Sun, C., Zhu, H., Clark, S. & Gouaux, E. Cryo-EM structures reveal native GABAA receptor assemblies and pharmacology. Nature 622, 195–201 (2023).

21. Kim, J. J. et al. Shared structural mechanisms of general anaesthetics and benzodiazepines. Nature 585, 303–308 (2020).

22. Laverty, D. et al. Cryo-EM structure of the human α1β3γ2 GABAA receptor in a lipid bilayer. Nature 565, 516–520 (2019).

23. Masiulis, S. et al. GABAA receptor signalling mechanisms revealed by structural pharmacology. Nature 565, 454–459 (2019).

24. Phulera, S. et al. Cryo-EM structure of the benzodiazepine-sensitive α1β1γ2S tri-heteromeric GABAA receptor in complex with GABA. eLife 7, (2018).

25. Zhu, S. et al. Structure of a human synaptic GABAA receptor. Nature 559, 67–72 (2018).

26. Zhu, S. et al. Structural and dynamic mechanisms of GABAA receptor modulators with opposing activities. Nat. Commun. 13, 4582 (2022).

27. Legesse, D. H. et al. Structural insights into opposing actions of neurosteroids on GABAA receptors. Nat. Commun. 14, 5091 (2023).

28. Chojnacka, W., Teng, J., Kim, J. J., Jensen, A. A. & Hibbs, R. E. Structural insights into GABAA receptor potentiation by Quaalude. Nat. Commun. 15, 5244 (2024).

29. Noviello, C. M., Kreye, J., Teng, J., Prüss, H. & Hibbs, R. E. Structural mechanisms of GABAA receptor autoimmune encephalitis. Cell 185, 2469–2477.e13 (2022).

30. Kash, T. L., Jenkins, A., Kelley, J. C., Trudell, J. R. & Harrison, N. L. Coupling of agonist binding to channel gating in the GABA(A) receptor. Nature 421, 272–275 (2003).

31. Kash, T. L., Trudell, J. R. & Harrison, N. L. Structural elements involved in activation of the gamma-aminobutyric acid type A (GABAA) receptor. Biochem. Soc. Trans. 32, 540– 546 (2004).

32. O’Shea, S. M. & Harrison, N. L. Arg-274 and Leu-277 of the gamma-aminobutyric acid type A receptor alpha 2 subunit define agonist efficacy and potency. J. Biol. Chem. 275, 22764–22768 (2000).

33. Hales, T. G. et al. An asymmetric contribution to gamma-aminobutyric type A receptor function of a conserved lysine within TM2-3 of alpha1, beta2, and gamma2 subunits. J. Biol. Chem. 281, 17034–17043 (2006).

34. Nemecz, Á., Prevost, M. S., Menny, A. & Corringer, P.-J. Emerging Molecular Mechanisms of Signal Transduction in Pentameric Ligand-Gated Ion Channels. Neuron 90, 452–470 (2016).

35. Kim, J. J. & Hibbs, R. E. Direct Structural Insights into GABAA Receptor Pharmacology. Trends Biochem. Sci. 46, 502–517 (2021).

36. Nors, J. W., Gupta, S. & Goldschen-Ohm, M. P. A critical residue in the α1M2-M3 linker regulating mammalian GABAA receptor pore gating by diazepam. eLife 10, (2021).

37. Nors, J. W., Endres, Z. & Goldschen-Ohm, M. P. GABAA receptor subunit M2-M3 linkers have asymmetric roles in pore gating and diazepam modulation. Biophys. J. 123, 2085–2096 (2024).

38. Scheller, M. & Forman, S. A. Coupled and uncoupled gating and desensitization effects by pore domain mutations in GABA(A) receptors. J. Neurosci. 22, 8411–8421 (2002).

39. Keramidas, A. & Harrison, N. L. The activation mechanism of alpha1beta2gamma2S and alpha3beta3gamma2S GABAA receptors. J. Gen. Physiol. 135, 59–75 (2010).

40. Mortensen, M., Wafford, K. A., Wingrove, P. & Ebert, B. Pharmacology of GABA(A) receptors exhibiting different levels of spontaneous activity. Eur. J. Pharmacol. 476, 17– 24 (2003).

41. Horenstein, J., Riegelhaupt, P. & Akabas, M. H. Differential protein mobility of the gamma-aminobutyric acid, type A, receptor alpha and beta subunit channel-lining segments. J. Biol. Chem. 280, 1573–1581 (2005).

42. Horenstein, J., Wagner, D. A., Czajkowski, C. & Akabas, M. H. Protein mobility and GABA-induced conformational changes in GABA(A) receptor pore-lining M2 segment. Nat. Neurosci. 4, 477–485 (2001).

43. Pettersen, E. F. et al. UCSF ChimeraX: structure visualization for researchers, educators, and developers. Protein Sci. 30, 70–82 (2021).

44. Herschlag, D. & Pinney, M. M. Hydrogen Bonds: Simple after All? Biochemistry 57, 3338–3352 (2018).

45. Hubbard, R. E. & Kamran Haider, M. Hydrogen bonds in proteins: role and strength. in *Encyclopedia of life sciences* (ed. John Wiley & Sons, Ltd) (John Wiley & Sons, Ltd, 2001). doi:10.1002/9780470015902.a0003011.pub2.

## Additional References for Methods

46. Venkatachalan, S. P. et al. Optimized expression vector for ion channel studies in Xenopus oocytes and mammalian cells using alfalfa mosaic virus. Pflugers Arch. 454, 155–163 (2007).

47. Boileau, A. J., Baur, R., Sharkey, L. M., Sigel, E. & Czajkowski, C. The relative amount of cRNA coding for gamma2 subunits affects stimulation by benzodiazepines in GABA(A) receptors expressed in Xenopus oocytes. Neuropharmacology 43, 695–700 (2002).

48. Jorgensen, W. L., Chandrasekhar, J., Madura, J. D., Impey, R. W. & Klein, M. L. Comparison of simple potential functions for simulating liquid water. J. Chem. Phys. 79, 926 (1983).

49. Jo, S., Kim, T., Iyer, V. G. & Im, W. CHARMM-GUI: a web-based graphical user interface for CHARMM. J. Comput. Chem. 29, 1859–1865 (2008).

50. Best, R. B. et al. Optimization of the additive CHARMM all-atom protein force field targeting improved sampling of the backbone φ, ψ and side-chain χ(1) and χ(2) dihedral angles. J. Chem. Theory Comput. 8, 3257–3273 (2012).

51. Vanommeslaeghe, K. et al. CHARMM general force field: A force field for drug-like molecules compatible with the CHARMM all-atom additive biological force fields. J. Comput. Chem. 31, 671–690 (2010).

52. Abraham, M. J. et al. GROMACS: High performance molecular simulations through multi-level parallelism from laptops to supercomputers. SoftwareX 1–2, 19–25 (2015).

53. Bussi, G., Donadio, D. & Parrinello, M. Canonical sampling through velocity rescaling. J. Chem. Phys. 126, 014101 (2007).

54. Parrinello, M. & Rahman, A. Crystal structure and pair potentials: A molecular-dynamics study. Phys. Rev. Lett 45, 1196–1199 (1980).

55. Essmann, U. et al. A smooth particle mesh Ewald method. J. Chem. Phys. 103, 8577 (1995).

56. Hess, B., Bekker, H., Berendsen, H. J. C. & Fraaije, J. G. E. M. LINCS: A linear constraint solver for molecular simulations. Journal of Computational Chemistry (1997).

57. Michaud-Agrawal, N., Denning, E. J., Woolf, T. B. & Beckstein, O. MDAnalysis: a toolkit for the analysis of molecular dynamics simulations. J. Comput. Chem. 32, 2319– 2327 (2011).

58. Zhuang, Y. MD Simulations of a1b2g2 GABA-A receptor. Zenodo (2023) doi:10.5281/zenodo.8142630.

